# digimouth: a Python package for modeling aroma release in food science

**DOI:** 10.64898/2026.09.30.755647

**Authors:** C. Peltier, O. Mourabit, K. Coulibaly, I. Andriot, I.C. Trelea

**Affiliations:** Université Bourgogne Europe, Institut Agro, CNRS, INRAE, UMR CSGA, 21000 Dijon, France; Probe Research Infrastructure, ChemoSens facility, CNRS, INRAE, 21000 Dijon, France; Université Paris-Saclay, INRAE, AgroParisTech, UMR SayFood, F-91120, Palaiseau, France

## Abstract

The modeling of aroma release in food science is a complex, multidisciplinary field integrating mass transfer theory and principles of physical chemistry. While mechanistic models have long been used to understand volatile release, their practical adoption has been hindered because existing implementations are often private, “in-house” tools that lack reproducibility and extensibility. This paper introduces digimouth, an open-source Python framework designed to provide a unified, modular, and reproducible environment for simulating and fitting aroma release kinetics.

A key innovation of digimouth is its handling of events and stages. Events represent discrete, instantaneous changes in the system (such as opening a bottle or swallowing), while stages allow for multi-phase simulations where experimental conditions or physical dynamics change over time.

To demonstrate the tool’s utility, the authors implemented established *in vitro* convection models and performed sensitivity analyses on parameters such as air-water partition coefficients and mass transfer coefficients. The package also includes an optimization module based on the least-squares method to fit parameters to experimental data. Validation against real-time data from Proton Transfer Reaction–Time of Flight– Mass Spectrometry (PTR-ToF-MS) shows a satisfactory fit, with an average error of approximately 2%.

Ultimately, digimouth facilitates robust model development and hypothesis testing, offering the food science community a collaborative tool to better understand the mechanisms of flavor release.

## 1. Introduction and context

The modeling of aroma release in food systems is a multidisciplinary challenge that integrates physicochemical principles, mass transfer theory, and dynamic processing conditions. Early modeling efforts primarily relied on *in vitro* approaches to elucidate the fundamental mechanisms governing volatile release following a disturbance of equilibrium, such as exposure of a food product to air. These foundational studies identified key controlling parameters, including air–product partition coefficients and mass transfer coefficients in both liquid and gas phases (Marin et al., 1999), thereby establishing the mechanistic basis for subsequent modeling developments.

Building on these foundations, later models extended mass transfer frameworks to more complex food matrices, such as emulsions and semi-solid systems (Andriot et al., 2000; Harrison and Hills, 1997), where interactions between aroma compounds and food constituents can significantly influence release kinetics. In these systems, external mass transfer at the product–air interface is often identified as the rate-limiting step, although chemical binding and dissociation phenomena may also contribute under certain conditions.

Despite the conceptual and predictive power of these *in vitro* mechanistic models, their practical adoption remains limited. Most of the existing implementations have been developed as in-house research tools and are not publicly available, reusable, or easily extensible by external users. As a result, the broader community lacks access to a unified computational framework capable of integrating these established modeling approaches, hindering reproducibility, model comparison, and further methodological development.

This work introduces a modular simulation framework for aroma-release dynamics that unifies ODE (ordinary differential equations) -based models, event-driven state transitions, multi-stage experimental protocols, and parameter estimation within a single computational environment. The capabilities of the framework are demonstrated through simulations reproducing key aroma-release mechanisms described in the literature. Its predictive performance is subsequently evaluated by comparing model outputs with experimental *in vitro* data.

## 2. Material and methods

### 2.1. Implemented framework

#### 2.1.1. Design of the tool

The tool presented in this paper is a Python simulation framework (digimouth) designed to simulate aroma-release dynamics *in vitro* and, potentially, *in vivo*. It is organized into four main modules. The first (Module 1) explicitly implements existing models, such as those developed by Marin et al. (1999). The second computes the system dynamics based on a given model, parameter values, initial conditions, and optional discrete events. The third provides graphical tools for visualizing simulation and optimization results. Finally, the fourth performs parameter estimation through optimization procedures. While the second module is generic and handles the simulation of system dynamics independently of the underlying model, the first one provides implementations of specific published or newly developed models. Together, these four modules provide a unified framework for model definition, simulation of complex experimental protocols, parameter estimation, and result visualization. They are presented in the next sections and illustrated in Figure 1 (details in Appendix A).

**Figure 1:**
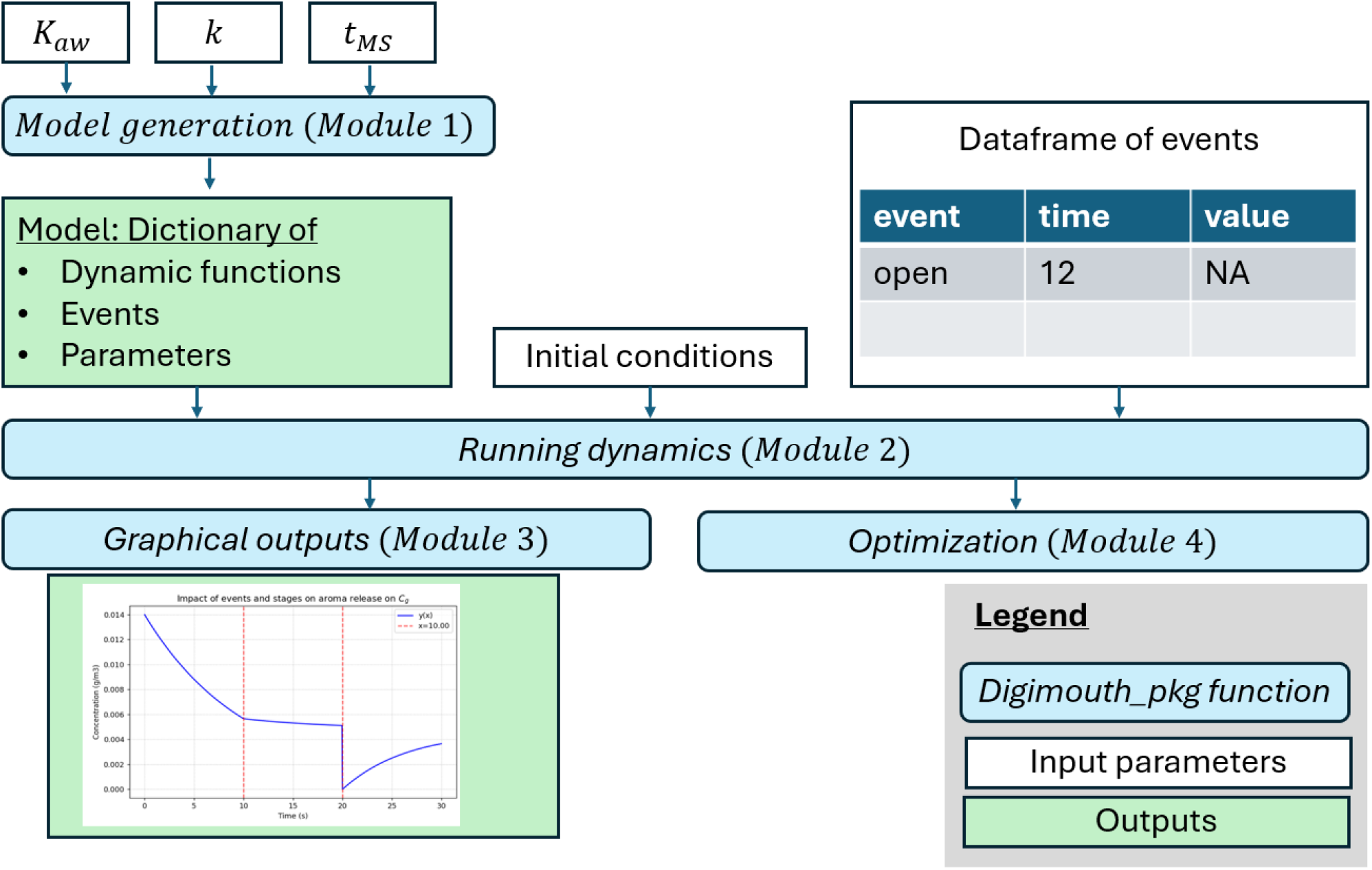
Pipeline of the tool of aroma release simulation and optimization

#### 2.1.4. Implemented models (Module 1)

As a starting point and a basic use case, the package implements the *in vitro* convection models described by Marin et al. (1999) and aims to be completed by more complex models (e. g. *in vivo*).

The *in vitro* model describes the release of volatile compounds from a liquid matrix into the gas phase and their subsequent detection by a measuring instrument. The system is represented as a closed headspace setup initially at thermodynamic equilibrium, followed by dynamic gas extraction by the analytical instrument. It is modeled using three well-mixed compartments corresponding to the liquid phase, the gaseous headspace, and a measurement compartment representing the PTR-MS tubing and chamber (Fig. 2).

**Figure 2:**
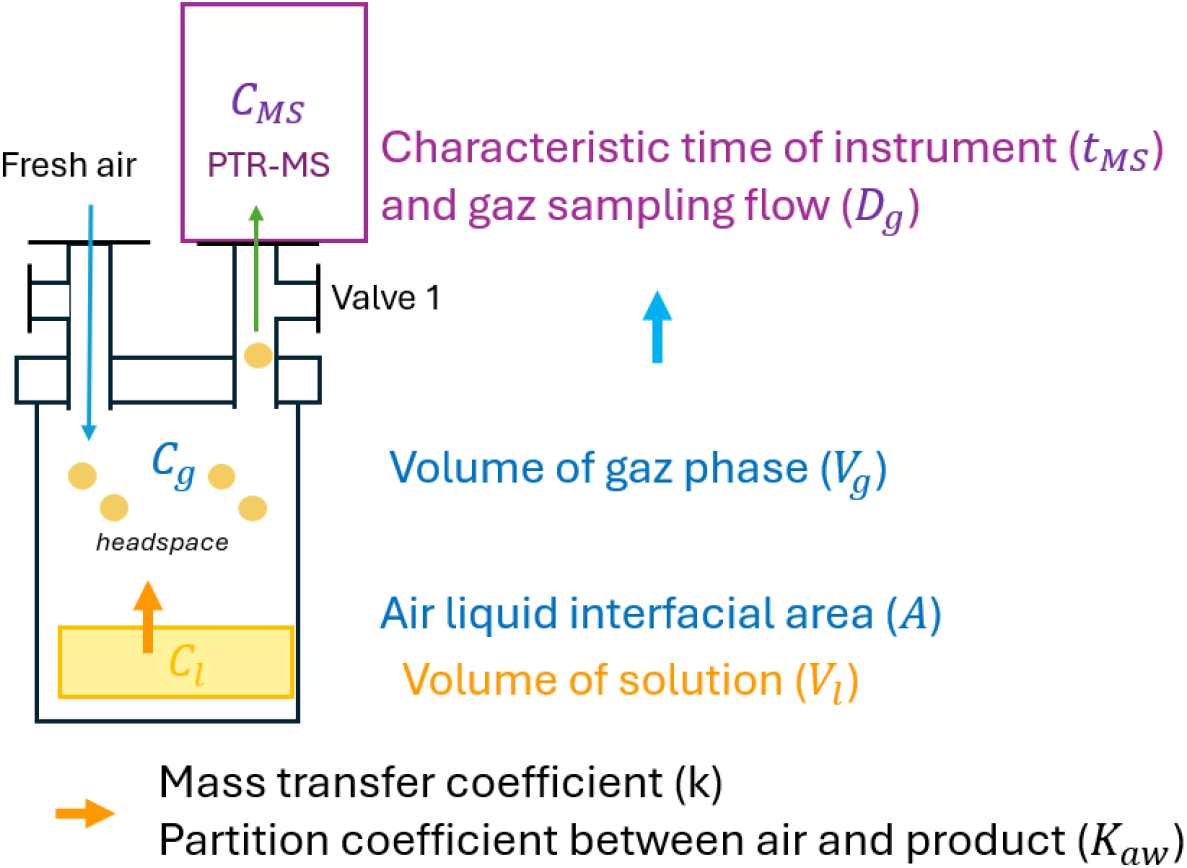
Experimental device. *C*_*l*_, *C*_*g*_ and *C*_*MS*_ are respectively the concentrations of the volatile compound in the liquid phase, gas headspace and PTR-MS compartments

Marin et al. (1999) established the following system of ordinary differential equations based on dynamic mass balances in the specified compartments:

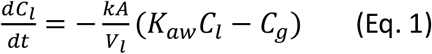

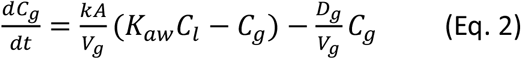

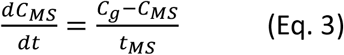

Eq. 1 describes the removal of the volatile compound from the liquid compartment towards the gaseous headspace (see Appendix B for parameter significance and units). The first term in Eq. 2,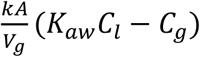 represents mass transfer across the air–liquid interface. Here, the driving force corresponds to the departure from thermodynamic equilibrium. The second term, 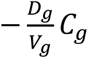 accounts for the continuous removal of headspace gas by the PTR-MS. The PTR-MS samples the headspace at a constant volumetric flow rate *D*_*g*_, representing a convective loss of volatile compounds from the gas phase. This term is essential to describe the progressive depletion of the headspace during measurement.

The PTR-MS does not instantaneously measure the air concentration. Instead, the signal is affected by tubing transport, instrumental response time, mixing and residence time. This is modeled as a first-order low-pass filter (Eq. (3)), where *t*_*MS*_ represents the characteristic response time of the measurement system.

#### 2.1.2. Resolution of dynamics (Module 2)

In the presented framework, a model is defined by:

- a set of state variables,
- their associated dynamics represented by ordinary differential equations (ODEs)
- their related parameters (that could be optimized)
- a collection of optional event functions.

These event functions constitute a distinctive feature of the package. They represent instantaneous modifications of the system state, such as very fast mass transfers, rapid volume changes, or resets of specific variables, occurring at predefined times and generating discontinuities that cannot be described by ODEs alone. This mechanism is particularly relevant for modeling phenomena that occur on timescales much shorter than that of the overall experiment. By explicitly accounting for such rapid transitions, the framework enables the simulation of complex experimental protocols involving abrupt physical or physiological changes.

The concept of stage is another distinctive feature of the package. A stage corresponds to a time interval during which a specific ‘model’ applies (i.e. set of equations, parameters and events). By combining successive stages, the framework can represent multiphase protocols involving changes in experimental conditions, such as container opening in *in vitro* conditions or swallowing the product in *in vivo* conditions.

Together, these features provide a flexible framework for robust event-driven and multi-stage ODE simulations. While particularly suited for applications in food science, such as aroma release, the framework remains broadly applicable to many dynamic modeling contexts. Events and stage transitions are managed through a restart strategy to ensure numerical stability and convergence of ODE solutions.

Numerical integration of ODE is performed using the LSODA algorithm from SciPy (Hindmarsh, 1983; Petzold, 1983), which automatically handles both stiff and non-stiff systems.

#### 2.1.3. Parameter optimization (Module 4)

The parameters used in the differential equations are not always known by the experimenter. Consequently, a module was implemented in order to optimize them according to the experimental data. Note that when a parameter has the same name in the different equations, it is considered as the same parameter and is optimized as a single value (and not separately in the different compartments). The optimization method is based on the least square method.

When experiments are repeated, multiple curves can be included in the optimization procedure. A first approach consists of optimizing each curve independently and then averaging the estimated parameter values. Alternatively, the package implements a global optimization approach in which a single set of parameters is estimated from the concatenation of all experimental curves. An advantage of this approach is that it generally yields narrower confidence intervals than independent optimizations, as all the data are considered simultaneously. In this case, the residuals are computed by comparing the concatenated experimental curves with the corresponding concatenation of the model simulations replicated for each experimental curve. The optimization function also includes an optional normalization step (enabled by default), allowing the optimization to focus on the kinetics (i.e., the shape of the curves) rather than their amplitudes.

### 2.2. Package usage

#### 2.2.1. Baseline simulations

Simulations of the *in vitro* setting were performed to quantify the impact of the model parameters. The following baseline parameter values were used : *K*_*aw*_ = 7 · 10^−3^; *k* = 1.38 10^−4^ *m. s*^−1^; *C*_0_ = 2 *kg* · *m*^−3^; *A* = 1 · 10^−3^ *m*^2^; *D*_*g*_ = 2.63 · 10^−6^ *m*^3^. *s*^−1^; *V*_*l*_ = 2 10^−6^ *m*^3^; *V*_*g*_ = 28 · 10^−6^ *m*^3^; *t*_*MS*_ = 1*s*.

Then, the results of simulations were plotted for each compartment (in the product, in the headspace, in the PTR-MS instrument).

#### 2.2.2. Impact of parameters on aroma release kinetics

Starting from the baseline values described in section 2.2.1, one parameter among *K*_*aw*_, *k, D*_*g*_ and *t*_*MS*_ was varied at a time in order to assess its influence on the release kinetics over the time interval 0–100 s. For each parameter, three values were considered: half the baseline value, the baseline value, and twice the baseline value.

#### 2.2.3. Illustrating the use of events and multistage options

To illustrate the use of events and stages, we considered the following situation: we consider that in a first period (0, 10 *s*), *k* = 1.38 · 10^−4^ *m. s*^−1^ corresponding to a static solution and then from 10 to 30 *s, k* = 10 * 1.38 · 10^−4^ *m. s*^−1^ corresponding to a mechanically stirred liquid solution. Then, we consider that an event ‘opening the vial’ occurs at t=20 s. In this case, the event consists in setting *C*_*g*_ = 0 at the given time (t=20*s*) and assess the effect on the results on the aroma release kinetics. Resulting simulated concentration of *C*_*g*_ were reported and plotted in this situation.

#### 2.2.4. Optimization of parameters with real dataset

Experimental data was collected to compare the results of simulations to reality. *In vitro* data was extracted of the experiments presented in Andriot et al. (2026). We consider there 2 mL of a solution of isoamylacetate in water (2ppm) into a 50 mL vial. The headspace was equilibrated for 5 minutes. Headspace analyses were performed with a PTR-ToF-MS (Proton Transfer Reaction – Time of Flight -Mass Spectrometry, PTR-ToF-8000, Ionicon Analytik GmbH, Innsbruck, Austria), an instrumental device that allows aroma release to be quantified along time, with a flow Dg=2.63 (cm^3^/s). Each experiment lasted 3 minutes: 30 seconds for the blank (i.e., ambient air), 2 minutes for the vial headspace, and 30 seconds for the final blank. Only the period of aroma release during 2 minutes was analyzed here. Three replicates were conducted. We present the use of parameter optimization on three in vitro experiments for quantifying the mass transfer coefficient (*k*) and the response time of PTR-MS (*t*_*MS*_). The partition coefficient at 25°C (*K*_*aw*_) is fixed at *K*_*aw*_ = 2.3 · 10^−2^, a value obtained from separate GC experiments, using the phase ratio method described inTromelin et al. (2012).

The parameters *t*_*MS*_ and *k* were optimized. The optimization bounds were (1 · 10^−6^, 1 · 10^−3^) for *k*, in *m. s*^−1^ and (1 · 10^−3^, 2) for *t*_*MS*_ in *s*.

The three experimental curves were first modeled separately, then the three curves together with a unique set or parameters. Each time, the simulated and the experimental data are plotted and optimal parameters and RMSE are given.

### 2.3. Computational aspects and availability

The package was developed and tested using Python 3.12. Each implemented function is associated with a dedicated test function including examples that can be run by the user, facilitating debugging, validation, and long-term maintainability of the codebase (Appendix B). The package is freely available on github (https://github.com/Chemosens/digimouth).

## 3. Results

### 3.1. Baseline simulations

Figure 3 shows the behavior of the three *in vitro* concentrations in baseline conditions. As expected, the concentration in the solution slightly decreases from an initial value of 2 *kg* · *m*^−3^. The decrease represents less than 5% at the experiment time scale (100 s). The concentration in the gas phase (headspace) starts at its equilibrium value *C*_0_*K*_*aw*_ then decreases roughly following a decaying exponential with a characteristic time *V*_*g*_/*D*_*g*_, corresponding to the headspace wipeout. The final pseudo-plateau represents a dynamic equilibrium between volatile supply from the solution and extraction by the gas flow. It is slightly decreasing, at the time scale of the solution depletion. Finally, the concentration measured by the PTR-MS exhibits a slightly lower peak than in the headspace, illustrating the low-pass filter effect of the measuring instrument. Its long right-hand tail essentially follows the concentration in the headspace.

**Figure 2:**
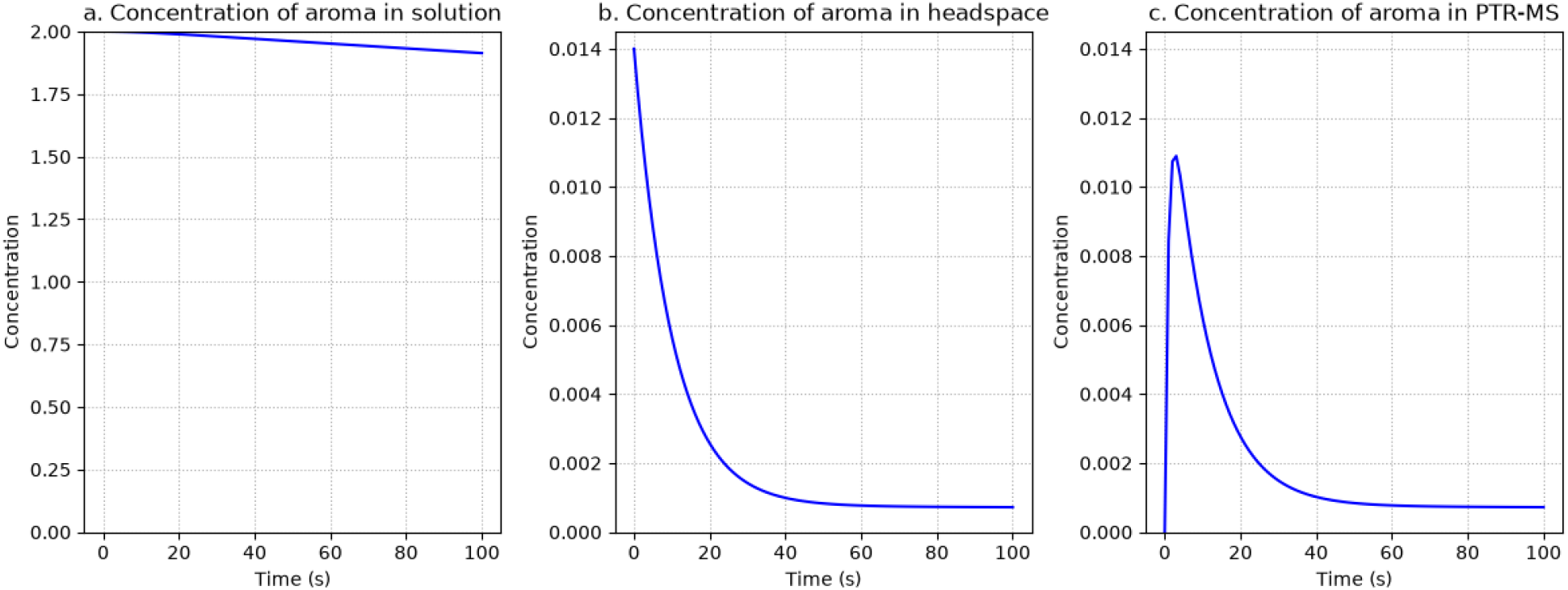
Concentration of aroma in solution (a), headspace (b), and PTR-MS (c)

**Figure 3:**
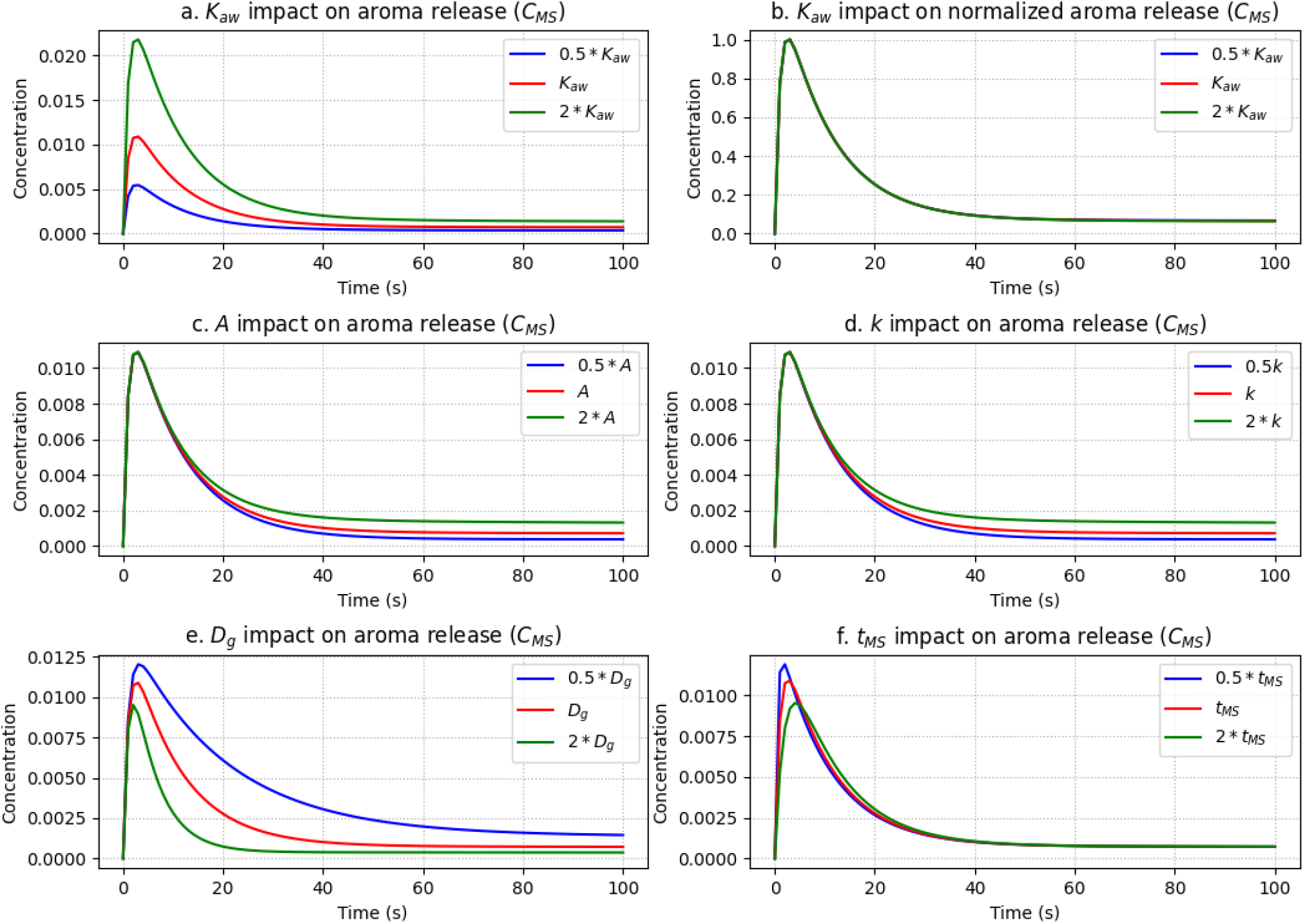
Impact of model parameters on the modeled kinetics of aroma release (*C*_*MS*_) for a 0-100 *s* simulated experiment.

### 3.2. Impact of parameters on aroma release kinetics

Figure 3 shows that all curves of *C*_*MS*_ vs time exhibit a peak-shaped profile. Increasing *K*_*aw*_ mainly affects the peak height: higher values of *K*_*aw*_ lead to higher peaks, since the initial headspace concentration is that of thermodynamic equilibrium with the liquid (Fig. 3a). Note that the shape of the normalized curve is not impacted by *K*_*aw*_(Fig. 3b). Regarding the *A* or *k* parameters, they appear to change the asymptotic value of the aroma concentration: higher values lead to higher asymptotic values, as expected, since the pseudo-plateau represents a dynamic equilibrium between aroma supply and removal (Fig 3c and 3d). An increase in *D*_*g*_ affects the width of the peak: higher *D*_*g*_ values result in a narrower peak, in agreement with the observation that the characteristic time of the removal is *V*_*g*_/*D*_*g*_. Also, higher *D*_*g*_ results in a lower plateau, still in agreement with the dynamic equilibrium remark. This is consistent with the interpretation that a high *D*_*g*_ corresponds to stronger aspiration by PTR-MS, leading to faster and stronger volatile removal from the headspace. Finally, higher *t*_*MS*_ values shift the curve to the right and reduce the peak height, illustrating the low-pass filter effect.

### 3.3. Illustrating the use of events and multistage options

Figure 4 illustrates the impact of discrete events and the multistage simulation.

**Figure 4:**
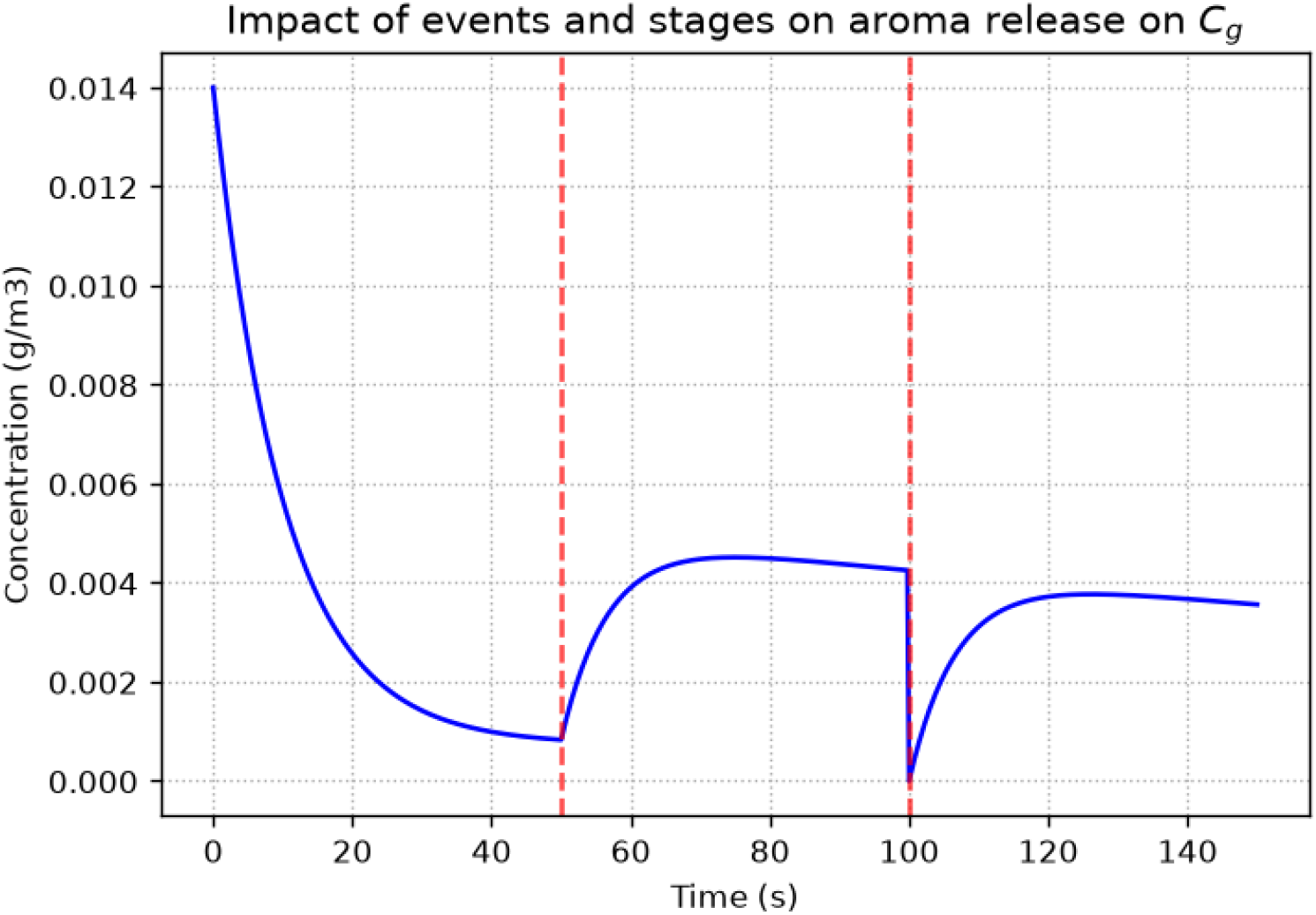
Impact of multistage and event conditions on aroma release kinetics

Regarding the multistage simulation, the mass transfer coefficient (*k*) was increased tenfold at (t = 50 s) to illustrate enhanced stirring of the solution. This change induces a transient increase in the gas phase aroma concentration toward a higher pseudo-asymptotic concentration than before the stirring event.

The occurrence of the **‘opening the bottle’** event at (t = 100 s) has a marked effect on the aroma release kinetics. The gas-phase concentration is instantaneously reset to zero, after which aroma release resumes, increasing rapidly before reaching a maximum and then gradually decreasing toward a pseudo-asymptotic concentration as the aroma is progressively depleted from the liquid reservoir.

### 3.4. Illustration of optimization with experimental data

The optimization of two model parameters (*k* and *t*_*MS*_) based on experimental data yields the results presented in Figure 6 and Table 1. In this case, the RMSE values are lower than 0.02 on a scale of 1 (after the maximum peak value was normalized to 1), corresponding to an average error of less than 2%. Overall, the fit is quite satisfactory.

**Table 1.** Estimated parameters and RMSE for the optimized model based on three samples.

| | $k$ | Confidence interval ($k$) | $t_{MS}$ | Confidence interval ($t_{MS}$) | RMSE |
| --- | --- | --- | --- | --- | --- |
| expe_1 | $2.09 \cdot 10^{-4}$ | $[2.04, 2.14] \cdot 10^{-4}$ | 0.94 | $[0.923, 0.962]$ | $1.92 \cdot 10^{-2}$ |
| expe_2 | $2.48 \cdot 10^{-4}$ | $[2.44, 2.52] \cdot 10^{-4}$ | 0.78 | $[0.781, 0.785]$ | $1.50 \cdot 10^{-2}$ |
| expe_3 | $2.42 \cdot 10^{-4}$ | $[2.4, 2.44] \cdot 10^{-4}$ | 0.88 | $[0.868, 0.884]$ | $0.84 \cdot 10^{-2}$ |
| expe_all | $2.32 \cdot 10^{-4}$ | $[2.29, 2.33] \cdot 10^{-4}$ | 0.88 | $[0.868, 0.887]$ | $1.58 \cdot 10^{-2}$ |
| Average of individual estimations | $2.27 \cdot 10^{-4}$ | $[1.81, 2.85] \cdot 10^{-4}$ | 0.87 | $[0.666, 1.07]$ | $1.42 \cdot 10^{-2}$ |

**Figure 6:**
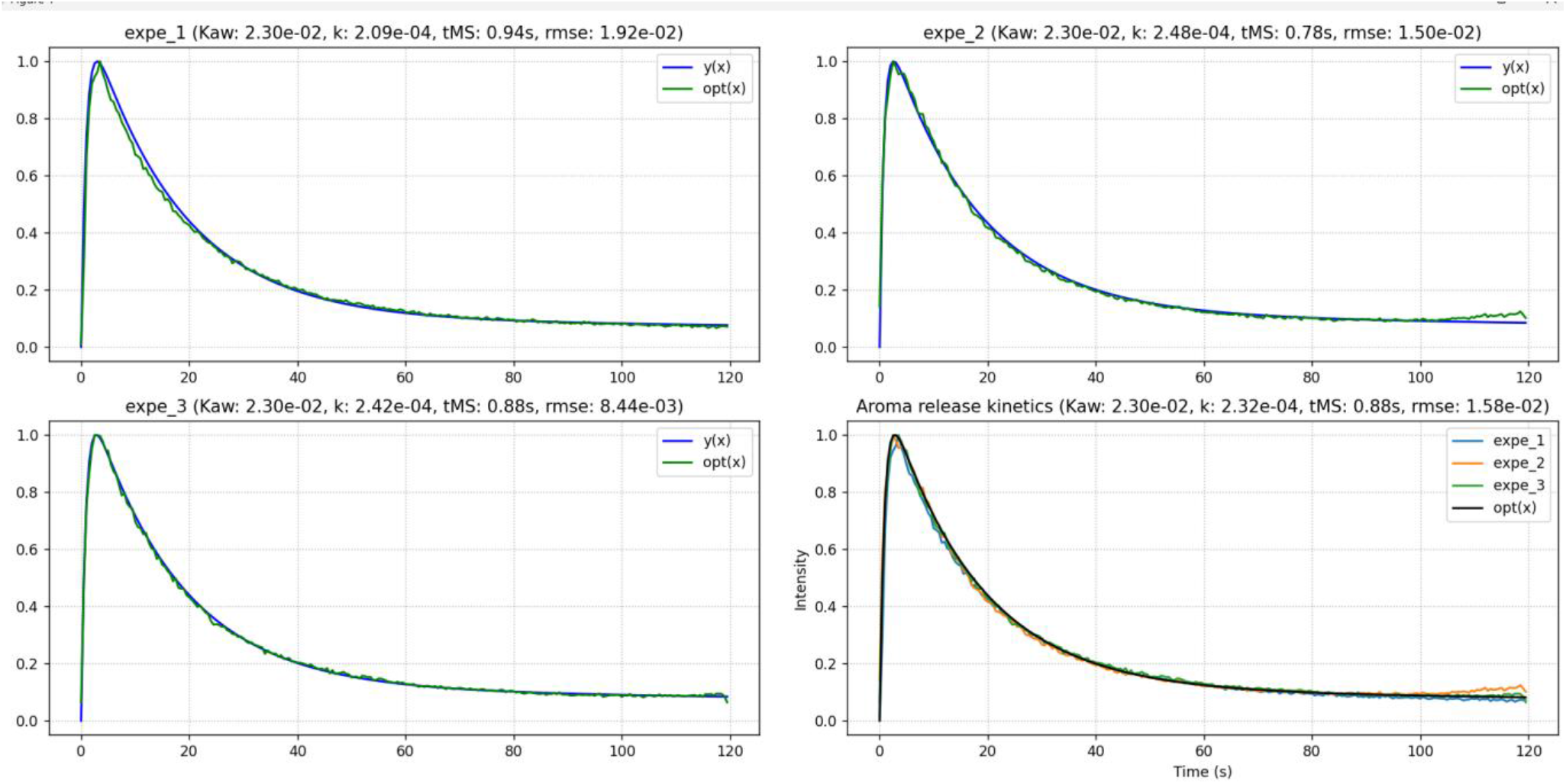
*In vitro* experimentation for the simultaneous optimization of *t*_*MS*_ and *k*. For expe_1, expe_2, expe_3, the experimental data is in green and the optimal simulated release is in blue. For the total aroma release kinetics, the three experimental curves are in blue, orange and green and the optimal release is in black.

However, the optimal parameter values differ for the individual experiments.

The *k* and *t*_*MS*_ parameters obtained from the global optimization are close to the average values of the three individual optimizations. However, the confidence intervals for the averaged parameters are much smaller (more than 20 times narrower) with the global optimization than with individual estimations.

## 4. Discussion

### 4.1. Interest of the package for *in vitro* analysis of aroma release

The use of *in vitro* data to model aroma release is of considerable interest, although it may introduce biases that are not always explicitly accounted for. In particular, repeated opening and closing of sample containers during pipette sampling are generally neither monitored nor incorporated into the modeling framework. Such handling may lead to non-negligible volatile losses, especially when containers remain open for extended periods or when highly volatile compounds are studied. As a result, the effective analyte concentration may progressively change over time, potentially affecting the accuracy and reproducibility of aroma release measurements and simulations.

The present package enables the explicit representation of repeated product manipulations and event-driven concentration changes during experimental workflows. In the present study, the implemented model describes a liquid solution governed by convection dynamics. However, the framework is sufficiently flexible to incorporate more complex physicochemical processes, such as dissolution and/or thermal degradation of food matrices during *in vitro* heating experiments. This would allow the simulation of coupled solid–liquid–gas transfer phenomena, possibly with chemical reactions, and more realistic dynamic food transformation scenarios.

### 4.2. Limits of the current models and their related experimental data

In this study, the data were normalized to focus solely on the kinetics. However, the amplitude of the signal is also of interest. The relationship between the actual concentration and the signal recorded by the PTR-MS device is not straightforward. Further work could aim to quantify the volatile concentration in the gas stream.

At present, the optimization framework is restricted to single-stage simulations, corresponding to the estimation of parameters for one model over a single continuous time interval. Extending the optimization procedure to multi-stage protocols remains possible but introduces additional complexity. In particular, stages may differ in duration, while some parameters can either remain shared across stages or vary independently between them. Such configurations require more advanced optimization strategies and parameter management. Consequently, users are currently encouraged to optimize each stage separately before combining them into a complete multi-stage simulation. The implementation of global multi-stage optimization procedures constitutes a perspective for future versions of the package.

### 4.3. Perspectives

Future perspectives include expanding the modeling framework to account for more complex product compositions, such as systems containing multiple aroma compounds as well as other constituents (e.g., sugars and fats). Interactions between these molecules could then be explicitly incorporated into the model to better reflect realistic food matrices.

Another promising direction is the integration of models describing solid food breakdown during oral processing i.e. *in vivo*, with a stronger emphasis on physical mechanisms. Such approaches could rely on advanced simulation tools (e.g., SOFA, finite element methods) to quantify structural changes during mastication and bolus formation. Indeed, SOFA provides a Python API and could therefore be called directly from the package. The collaborative nature of the package enables researchers in the field to test and integrate their own models, thereby fostering collective progress toward a better understanding of the underlying mechanisms.

## 5. Conclusions

The development of digimouth provides a concrete response to the food science community’s need for a unified, modular, and reproducible modeling framework for aroma release. By utilizing a flexible, dictionary-based ordinary differential equation (ODE) engine, this Python package allows researchers to simulate complex experimental protocols through the management of discrete events and successive stages.

Validation of the framework using *in vitro* convection models demonstrated its accuracy, with a satisfactory fit to Proton Transfer Reaction – Mass Spectrometry (PTR-MS) data showing an average error of approximately 2%. Beyond simple simulation, digimouth integrates a robust optimization module that enables the estimation of essential physicochemical parameters, such as mass transfer coefficients (*k*) and partition coefficients (*Kaw*), directly from experimental data.

A major strength of this tool is its open-source nature, which addresses the limitations of private “in-house” tools and encourages collaboration and model comparison across different laboratories. By facilitating hypothesis testing and the development of robust models, digimouth serves as a structuring tool for understanding the mechanisms of volatile compound release. Future perspectives for digimouth are extensive. They include expanding the framework to account for more complex food matrices (containing sugars or fats) and transitioning toward in vivo modeling.

## Acknowledgements

The research leading to these results has received funding under grant (24.7 DIGIMOUTH) from Carnot Qualiment© (DOI: 10.17180/h5gd-gk88) supported by Agence Nationale de la Recherche (20 CARN 0026). This work was performed thanks to CALIS network’s resources (doi: 10.15454/9PGN-W156).

## Appendix A : documentation of digimouth

The **digimouth** toolkit is a modular and reproducible Python package for modelling, simulating, and fitting the oral release and transport of volatile compounds. It is organized into the following modules:

- **existing_models.py**: contains implementations of *in vitro* models from the literature (Module 1)
- **dynamic_model.py**: implements a flexible dictionary-based ODE engine together with the concepts of **events** and **stages** for simulating complex dynamic systems. The generate_model function enables users to build models from scratch, while run_model executes a model simulation. The run_stages function allows sequential execution of multiple models (Module 2)
- **graphical_methods.py**: includes functions for visualization and graphical analysis of simulation and fitting results. (Module 3)
- **optimization.py**: provides tools for fitting models to experimental data.(Module 4)

### A.1. Model class (existing_model.py)

The Model class is the core object used in digimouth to represent a dynamic model. It encapsulates all the information required to define a system of ordinary differential equations (ODEs), including the model parameters, the differential equations, and optional event functions. This unified representation facilitates model construction, simulation, optimization, and reuse throughout the package.

A Model object contains the following attributes:

- params_dict: a nested dictionary containing the model parameters. Parameters are organized by state variable, allowing each equation to access its own set of parameters.
- funcs_dict: a dictionary associating each state variable with the function defining its time derivative.
- event_funcs (optional): a dictionary of event functions indexed by event name. Events represent instantaneous modifications of the system during a simulation, such as opening a bottle, swallowing, or changing a model parameter.
- name (optional): a descriptive name for the model.

Initial conditions are not stored within the Model object. Instead, they are provided separately when calling the simulation functions. This design allows the same model structure to be simulated under different initial conditions without redefining the model.

The Model class is a lightweight container and does not perform numerical integration itself. Simulations are carried out using the run_model() function, while models can either be constructed manually or generated automatically using the generate_model() helper function.

### A.2. Core simulation functions (dynamic_model.py)

The numerical engine of digimouth relies on a small set of generic functions that allow users to define, simulate, and analyze dynamic models independently of their mathematical formulation.

#### generic_model()

The generic_model() function evaluates the right-hand side of a system of ordinary differential equations (ODEs). State variables, differential equations, and parameters are provided as dictionaries, making the implementation independent of the number and names of variables.

Before evaluating the derivatives, the function verifies the consistency between the state variables, the differential equations, and the associated parameter dictionaries. It also checks that the parameters supplied to each differential equation match the function signature, providing informative error messages when inconsistencies are detected.

The function returns the derivatives of all state variables in the same order as the state vector.

#### resolution_odeint_generic()

The resolution_odeint_generic() function numerically integrates a dynamic model using the odeint solver from the SciPy library. A Model object together with the initial conditions and the simulation time vector are provided as inputs.

Internally, the function converts the state vector used by the numerical solver into dictionaries compatible with generic_model(), allowing models to be defined symbolically while remaining compatible with standard numerical integration routines. The simulation results are returned as a pandas.DataFrame, with one column per state variable and an additional time column.

#### apply_event_func()

The apply_event_func() function applies an instantaneous event to the current state of the model. Events are identified by their type and associated with user-defined event functions stored in the model. Typical events include modifying parameter values, resetting state variables, transferring mass between compartments, or changing simulation conditions.

#### run_model()

The run_model() function performs the simulation of a single stage. The simulation interval is automatically divided into successive segments separated by events. Each segment is numerically integrated using resolution_odeint_generic(), after which the corresponding event is applied instantaneously before continuing the simulation.

This approach allows discontinuous changes in parameters or state variables while maintaining continuous numerical integration between events. The complete simulation is returned as a single continuous data frame.

#### run_stages()

The run_stages() function executes a complete experimental protocol composed of successive simulation stages. Each stage may use a different Model object, corresponding to different differential equations and parameter values. The final state of one stage automatically becomes the initial condition of the next stage, allowing complex protocols involving multiple physical or physiological processes to be represented naturally. Events occurring within each stage are handled independently, and the outputs of all stages are concatenated into a single continuous simulation. The *run_stages* function returns a single concatenated pandas.DataFrame containing the temporal evolution of all state variables across the entire protocol. This unified timeline can subsequently be used for visualization, quantitative analysis, parameter estimation, or comparison with experimental data.

### A.3. Optimization module (optimization.py)

The optimization.py module provides a generic framework for calibrating dynamic models against experimental observations. It is fully compatible with the Model objects defined in digimouth and can optimize any model expressed as a system of ordinary differential equations.

#### Residual computation

Parameter estimation relies on minimizing the residuals between simulated and experimental data. The calculate_residuals() function updates the model parameters according to the current optimization vector, runs a numerical simulation using resolution_odeint_generic(), and computes the difference between simulated and experimental observations. Simulations may optionally be normalized to facilitate comparisons between datasets with different signal amplitudes. The function also supports parameter-dependent initial conditions through user-defined dependency functions.

The calculate_residuals_multi() function extends this approach to multiple experimental datasets. Rather than fitting each experiment independently, residuals from all experiments are concatenated into a single residual vector, allowing simultaneous estimation of shared model parameters.

#### Parameter optimization

Parameter estimation is performed using the optim_param_dynamic() function, which relies on the nonlinear least-squares algorithm implemented in scipy.optimize.least_squares. The user specifies the parameters to optimize, while all remaining parameters remain fixed.

The function automatically detects parameters shared by several state equations. Such parameters are optimized only once and their optimized value is propagated to every equation in which they appear. This considerably simplifies the definition of compartment models in which several differential equations depend on common physical parameters.

The optimization routine returns the optimized parameter values together with the optimized model, the residual sum of squares (SSR), the root mean squared error (RMSE), and optional estimates of parameter uncertainty.

The optim_param_dynamic_multi() function extends this procedure to the simultaneous fitting of multiple experimental datasets, while optim_param_dynamic_multiphases() performs parameter estimation for models composed of successive simulation stages separated by discrete events.

#### Confidence intervals

The package provides and estimates parameter uncertainty. It computes confidence intervals from the covariance matrix estimated from the Jacobian of the least-squares solution, assuming locally quadratic behavior of the objective function. Standard errors and confidence intervals are obtained using the Student’s *t*-distribution.

#### Initial parameter search

To reduce the sensitivity of nonlinear optimization to the initial parameter values, the grid_search_initial_guess() function performs an exhaustive search over a user-defined parameter grid. Each parameter combination is evaluated by computing the corresponding residual cost, and the combination producing the lowest mean squared error is selected as the starting point for subsequent nonlinear optimization.

This preliminary search is particularly useful for highly nonlinear models exhibiting multiple local minima but may require a large number of model simulations if many model parameters are fitted.

#### Input flexibility

The optimization module supports several input formats. Experimental data can be supplied as single vectors, lists of experiments, or dictionaries indexed by experiment identifiers through the optim_param_dynamic_dict() wrapper. Consequently, the same optimization framework can be applied to individual experiments, replicated measurements, or complete experimental campaigns without modifying the model implementation.

### A.4. Visualization module (graphical_methods.py)

The graphical_methods.py module provides visualization functions for simulated and experimental data generated with digimouth. Its main function, plot_xy(), offers a flexible interface for displaying one or several curves while remaining independent of the model structure.

The function accepts single datasets, collections of datasets (lists or dictionaries), and automatically adapts the plotting routine to the input format. Experimental and simulated data can be displayed simultaneously using different graphical representations (lines, markers, or both), facilitating visual comparison between observations and model predictions.

Additional features include customizable colours, axis labels, titles, axis limits, and the possibility of highlighting specific time points through vertical reference lines, which are particularly useful for representing discrete events occurring during a simulation (e.g., bottle opening, stirring, or swallowing).

The function can either create a new figure or plot on an existing Matplotlib axis, making it suitable for both standalone figures and multi-panel layouts. Figures may be displayed interactively or directly exported to publication-quality image files with user-defined resolution.

The plotting utilities are designed to provide a generic visualization interface compatible with all simulation and optimization functions implemented in digimouth, independently of the underlying mathematical model.

## Appendix B: Summary of different symbols used in the paper

Table 1 summarizes the different symbols that are used in the *in vitro* model.

**Table 1.** in vitro parameters and their unit.

| Symbol | Description | Units |
| --- | --- | --- |
| $C_g$ | Aroma concentration in the gas phase (headspace) | $kg \cdot m^{-3}$ |
| $C_l$ | Aroma concentration in the liquid phase | $kg \cdot m^{-3}$ |
| $C_{MS}$ | Aroma concentration measured by PTR-MS | $kg \cdot m^{-3}$ |
| $V_g$ | Gas (headspace) volume | $m^3$ |
| $V_l$ | Liquid volume | $m^3$ |
| $A$ | Air–liquid interfacial area | $m^2$ |
| $k$ | Overall mass transfer coefficient | $m \cdot s^{-1}$ |
| $K_{aw}$ | Air–water partition coefficient | dimensionless |
| $D_g$ | Gas sampling flow rate of the PTR-MS | $m^3 \cdot s^{-1}$ |
| $t_{MS}$ | PTR-MS response time constant | $s$ |

## Appendix C: Code for reproducing simulations

The code for reproducing simulations of this paper is available in digimouth package (examples) in digimouth_protocole_simulation_clean.py (for simulations) and digimouth_optim_in_vitro.py for comparison with experimental data.

